# Uncovering 2-D toroidal representations in grid cell ensemble activity during 1-D behavior

**DOI:** 10.1101/2022.11.25.517966

**Authors:** Erik Hermansen, David A. Klindt, Benjamin A. Dunn

**Affiliations:** Department of Mathematical Sciences, NTNU, Trondheim, Norway

**Keywords:** grid cells, topological data analysis, neural population dynamics

## Abstract

Neuroscience is pushing toward studying the brain during naturalistic behaviors with open-ended tasks. Grid cells are a classic example, where free behavior was key to observing their characteristic spatial representations in two-dimensional environments [1]. In contrast, it has been difficult to identify grid cells and study their computations in more restrictive experiments, such as head-fixed wheel running [2–6]. Here, we challenge this view by showing that shifting the focus from single neurons to the population level changes the minimal experimental complexity required to study grid cell representations. Specifically, we combine the manifold approximation in UMAP [7] with persistent homology [8] to study the topology of the population activity. With these methods, we show that the population activity of grid cells covers a similar two-dimensional toroidal state space during wheel running as in open field foraging [9, 10], with and without a virtual reality setup. Trajectories on the torus correspond to single trial runs in virtual reality and changes in experimental conditions are reflected in the internal representation, while the toroidal representation undergoes occasional shifts in its alignment to the environment. These findings show that our method can uncover latent topologies that go beyond the complexity of the task, allowing us to investigate internal dynamics in simple experimental settings in which the analysis of grid cells has so far remained elusive.

## Introduction

Recent neuroscience research has pushed for the study of naturalistic open-ended behaviors [11], suggesting the experimental complexity must be sufficiently high to make novel insights into neural computations [12]. The question, however, is what the minimal complexity is in order to study the function and ethological relevance of specific neurons [13, 14]. Grid cells are a canonical example from the brain’s navigational system [15] where the hexagonal grid patterns in their responses relative to space was first observed when the animals were allowed to freely move in large open field (OF) arenas [1, 16]. In contrast, the spatial responses of grid cells in one-dimensional (1-D) environments and head-fixed recordings are not yet well understood [17–19]: In 1-D linear tracks, these are speculated to be cross-sections of the 2-D patterns [6, 20], although recent results suggest that grid cells in 1-D tracks are tuned to integrated distance [21, 22]. Casali et al. even argued for ‘the necessity of identifying grid cells from real open field environments’ [2], thus implying that 2-D OF arenas are the minimal experimental complexity required to fully understand grid cell computations.

However, the brain performs computations not merely at the single-neuron level but also in populations of neurons (or *neural ensembles*) [reviewed in 23– 25]. The internal representations encoded by neural ensembles often carry a distinct topology [26–29] which can be studied without reference to external covariates [30, 31]. The concept of understanding this topological structure is beginning to change the way we think about certain classes of neurons [32, 33], such as head direction cells where the 1-D ring structure of the network represents the encoding of an internal head direction system [34, 35]. In the case of grid cells, their population representation has been found to be *toroidal*, describing the two periodicities of the grid cell pattern in 2-D environments [10]. Therefore, in this study we asked whether focusing on the topology of grid cell representations would allow us to study their computations in experiments of lower complexity where it has so far been impossible to give a convincing classification of grid cell populations [2].

### Population analysis

A common pipeline in population analysis starts by reducing the dimensionality of the neural activity, e.g., using uniform manifold approximation and projection (UMAP) [7], t-distributed stochastic neighbor embedding (t-SNE) [36] or (deep) latent variable models [37–39], with the goal of extracting a low-dimensional representation of the population activity at each time point. However, dimensionality-reduction requires specifying a range of parameters [40, 41], and in particular, choosing a dimensionality for the embedding space, often chosen as 2- or 3-D for interpretability [42, 43] which allows visualization but potentially discards crucial information [44]. Topological data analysis offers an alternative approach by characterizing highdimensional point clouds through *persistent homology* (PH) [8, 45]. Specifically, we can compute a topological invariant called a barcode quantifying the presence of holes in representations of the data. This characterization helps to discern underlying topological structure, such as circular features [46, 47]. However, the method is sensitive to outliers (Extended Data Fig. 5b) and has a computational bottleneck, with polynomial growth of run-time and memory requirements in the number of points in the point cloud [48]. Therefore, preprocessing (noise removal and downsampling) of the data is necessary to uncover shape information from the barcodes [10, 34, 35]. Here, we propose uniform manifold approximation and persistent homology (UMAPH), which resolves these issues by applying persistent homology to the topological representation constructed in the first step of UMAP, thus eliminating the potentially problematic projection step of UMAP [44], while providing a topology-preserving [7] noise model for PH (Fig. 1a and Extended Data Fig. 5a, b).

**Fig. 1.**
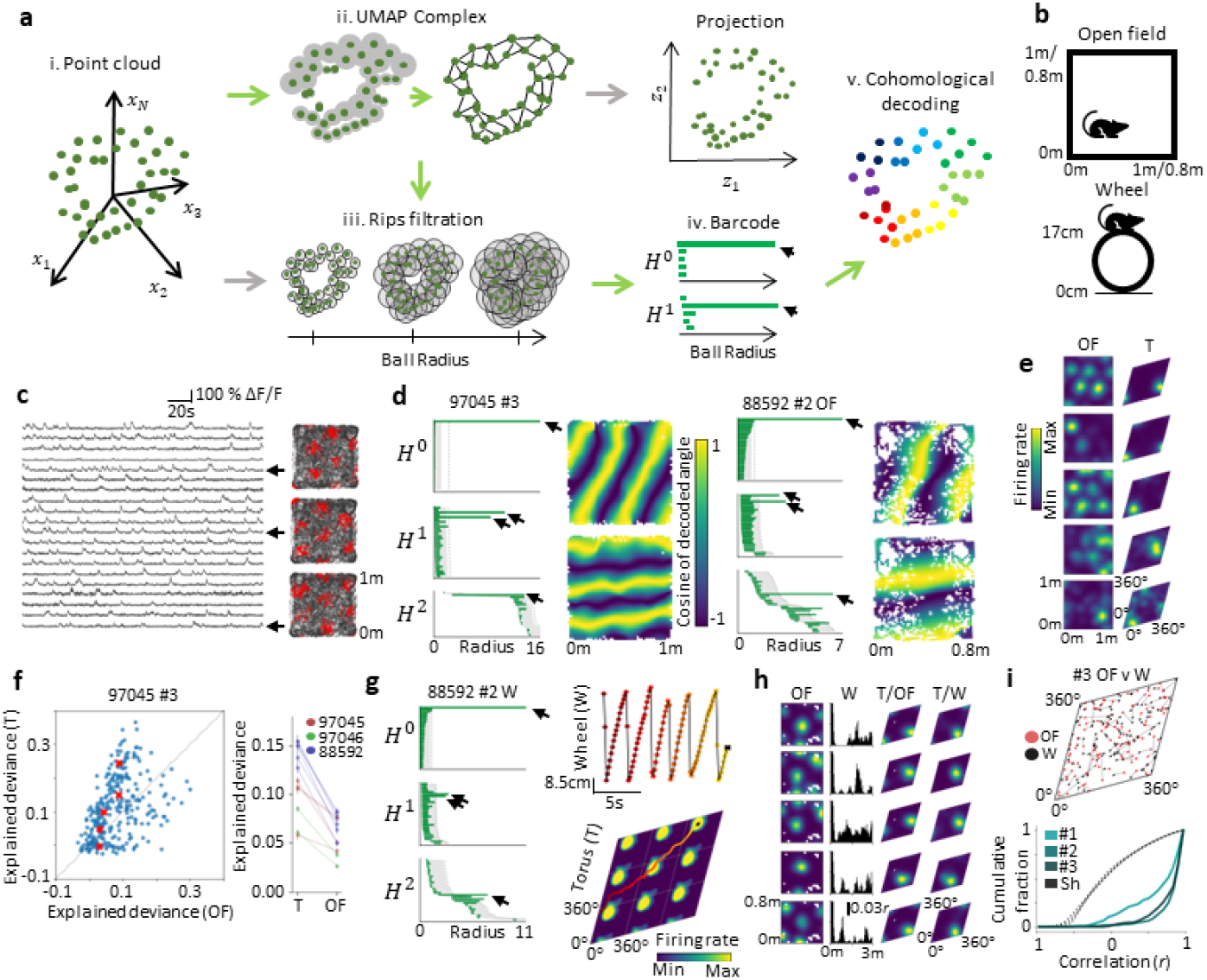
Toroidal representations in calcium imaging of mouse entorhinal cortex in 1- and 2-D environments. **a**. UMAPH (green arrows) combines aspects of UMAP (top arrows) and persistent homology (bottom arrows). Given a point cloud (green dots) in ambient space (i), UMAP computes a topological representation of the data (ii) based on local neighborhoods and approximates this in a low-dimensional projection of the data. Persistent homology constructs a sequence of combinatorial spaces from pairwise distances (iii) and tracks the evolution of *n*-dimensional holes (*H*^*n*^) in these spaces for increasing distances, displayed as bars in a barcode (iv). The long bars depict the most prominent features in the data (black arrows). 1-D *co*homology features allow for circular coordinatization (v). **b**. The mice ran in an OF arena (top) and on a Styrofoam wheel (bottom). **c**. 20 example calcium traces from the MEC (left). Coloring the spatial positions by heightened activity (red) of three example neurons reveals hexagonal grid patterns (right). **d**. Left in each block, barcodes from OF sessions in mice 97045 (day 3, *n* = 293 neurons) and 88592 (day 2, *n* = 160), indicating signature homology of a torus (one *H*^0^-bar, two *H*^1^-bars and one *H*^2^-bar). Only the 30 longest bars in each dimension are shown. Shaded gray shows 95th percentile and dotted line the maximum of the longest lifetimes of 100 shuffles. Right in each block, OF positions color-coded by mean decoded circular coordinates. **e**. Tuning of five example neurons (mouse 97045 day 3, see Fig. 7 for all neurons) to position in OF arena and on torus (T), sorted by explained deviance scores. **f**. Left, explained deviance scores for all recorded neurons in mouse 97045 day 3 describing goodness-of-fit for GLM models fitted to calcium events for T vs. OF positions as covariates. Red crosses mark the five cells shown in **e**. Right, mean value (*±*s.e.m.) for each OF (and object) session where toroidal topology was detected (Extended Data Fig. 1e, *n* ∈ [113, 637] neurons). **g**. Barcode from grid cell activity during wheel running of mouse 88592 day 2 indicates toroidal topology (left). Right, internal (bottom) and external (top) trajectory, color-coded by path progression, on the unwrapped torus during wheel running. Shading on the torus indicates mean population activity (indicated by color bar) at end of the path (black cross). **h**. Rate maps of top five neurons (mouse 88592 day 2), ranked according to toroidal explained deviance. From left: OF tuning; linearized wheel position autocorrelogram and tuning to torus inferred from corresponding sessions. Color-coding as in **e. i**. Top, toroidal tuning peaks of grid cell population in OF (red) and wheel (black) recordings (day 3). Lines connect peaks of the same neurons. Bottom, cumulative line plots of the correlation between rate maps from OF and wheel sessions for mouse 88592 days 1 – 3 (*n* = 126, 160 and 113 neurons). Dotted lines show shuffled distributions (*n* = 1000 shuffles).

### The UMAPH algorithm

We first compute geodesic distances of the data based on the assumption that a dataset is uniformly sampled from an underlying manifold (e.g., the grid cell torus), but is presented as a non-uniform distribution embedded into an ambient space with observational noise (e.g., neural population activity). This motivates obtaining the intrinsic distances by enforcing the volume of a ball containing a fixed number of points to be constant independent of which point in the dataset it circumscribes [7]. However, this creates local irregularities – the distance between two neighboring points will depend on the local neighborhood of each – so these are averaged to be symmetric (Fig. 1a.ii). UMAP then proceeds with an approximation of the same distances on a lower-dimensional point cloud. In UMAPH, we instead, apply PH to the high-dimensional data using the geodesic distances to compute the barcode of the manifold. PH first constructs a combinatorial description of the proximity between points. By forming balls of a common radius around each point, it lists the *sets of balls* with pairwise intersections. Increasing the radius forms a sequence of such descriptions, called a *filtration* (Fig. 1a.iii). The *n*-dimensional homology captures how unions of balls form *n*-dimensional holes. Thus, applying homology to the filtration describes the evolution of holes for increasing volumes, and the collection of radial intervals (or *bars*) when different holes appear and disappear is called a *barcode* (Fig. 1a.v). Optionally, in a last step, we perform cohomological coordinatization [49], to derive a low-dimensional description of the data (*decoding* the timevarying internal state [34], Fig. 1a.vi). Based on the selected 1-D holes in the barcodes, we compute circle-valued maps, assigning to each population vector angular coordinates corresponding to the circular features represented by the bars.

In summary, the intuition behind UMAPH is that we apply homology to a filtration constructed by forming balls containing the same number of points, but of potentially different sizes in the ambient space (Extended Data Fig. 5). This is advantageous because the metric is then locally adaptive to data density. Comparing the results when applied to head direction data previously studied with PH [30, 34, 35], gives an example of the benefit of this construction (Extended Data Fig. 6). With UMAPH, we found the resulting barcodes to be more interpretable with clear head direction rings maintained across states, even during slow-wave sleep where this structure was previously not recovered.

### Toroidal tuning of mouse entorhinal cells in open field calcium recordings

Zong et al. (2022, [50]) and Obenhaus et al. (2022, [51]) performed two-photon calcium imaging of the medial entorhinal cortex (MEC) in freely moving and head-fixed mice (Fig. 1b, c). In two mice (in 4/6 and 2/6 recording days), applying UMAPH to the activity of all recorded cells during OF firing resulted in barcodes with distinct toroidal signatures (one clear *H*^0^bar, two *H*^1^-bars and one *H*^2^-bar, all exceeding corresponding longest bars in 100 random perturbations of the data, Fig. 1d and Extended Data Fig. 1a). In a third mouse, we clustered the OF autocorrelograms to obtain a single grid cell ensemble (or *module* [9]), exhibiting similar toroidal expression in the barcode (in 4/7 OF recordings studied), also when an object was inserted (1 out of 3 object session recordings) (Fig. 1d and Extended Data Fig. 1a). The pair of circular coordinates attained from decoding of the two longest circular features in the barcode revealed stripe-like patterns in the OF corresponding to the two periodicities of the grid pattern (Fig. 1d and Extended Data Fig. 1a). Moreover, the internal dynamics displayed a clear relation to spatial movement (Video 1). We next studied each cell’s tuning to the torus by fitting a generalized linear model (GLM) to the calcium activity, comparing the performance when using either the toroidal positions or the recorded positions as covariates. The activity was better modeled by the torus than physical space (*P* ∈ [1.4 *·* 10^−34^, 0.0076], *W* ∈ [7694, 168511], Wilcoxon signed-rank test), and we observed that cells with the best fit had a clear grid pattern and single tuning fields on the torus (Fig. 1e, f and Extended Data Fig. 1e and 7).

### Linking toroidal grid cell tuning between open field and headfixed treadmill recordings

We next performed the same analysis of grid cell module activity while the animal was running on a wheel. A 2-D toroidal topology as in the OF sessions was seen (in 3 out of 4 wheel recordings), despite the 1-D nature of the task (Fig. 1g and Extended Data Fig. 1a). Decoding the positions on the torus revealed mostly unidirectional trajectories in line with the spatial behavior (Fig. 1g and Video 2). This shows that internal position integrates over the animal’s locomotion, even though the animal is not moving in space, supporting the idea of path integration. Each neuron had a single toroidal tuning field in the same location on the torus in OF foraging and wheel running (toroidal rate map correlations higher than shuffled comparisons, *n* = 1000 shuffles, *P <* 0.001, Fig. 1h, i and Extended Data Fig. 1c).

### Stable toroidal tuning of mouse grid cells in virtual reality with Neuropixels recordings

Campbell et al. (2021, [22]) used Neuropixels probes to record extracellular spikes in the MEC of head-fixed mice engaged in a virtual-reality task on a running wheel under different conditions: baseline, dark, gain (wheel slowed down with respect to virtual reality (VR)) and contrast (reduced visual contrast) sessions (Fig. 2a) [22]. We first studied one exemplary recording day (mouse I1 day 0417) with a good experimental yield (having 75 ‘distance’/’putative grid’ cells out of 225 cells recorded in the MEC using the classification given by Campbell et al.), before repeating our analysis for the remaining available data. Neuron ensembles were first clustered by their cross-correlation (maximized over *<* 0.9s time lags, Fig. 2b), thus avoiding any prior assumption about the spatial tuning of the studied neurons. We found two prominent clusters (henceforth termed ‘M1’ and ‘M2’, *n* = 44 and 41 neurons). Applying UMAPH to the firing rates of these clusters revealed barcodes with four long-lived bars (one *H*^0^-, two *H*^1^- and one *H*^2^-bar) indicating toroidal structure (*P <* 0.01 in all conditions, except M2 baseline: *P* = 0.06, Fig. 2c). Decoding revealed primarily unidirectional trajectories on the toroidal manifold (Fig. 2d). The activity of most cells in both M1 and M2 was confined to a specific location on the toroidal surface (Fig. 2e) and each neuron’s preferred location on the torus was preserved across the experimental conditions (*P <* 0.001 for toroidal rate map correlations compared to *n* = 1000 shuffles, Fig. 2e and Extended Data Fig. 8), consistent with grid cell populations in rats [10]. Lastly, to allow for unbiased comparison with the VR spatial track, we used the toroidal description found during the dark session to decode baseline trials and found the toroidal coordinates to have higher explained deviance than the VR positions during baseline trials (M1: *P* = 3.8 *·* 10^−9^, *W* = 990; M2: *P* = 1.2 *·* 10^−8^, *W* = 861, Wilcoxon signed-rank test; Fig. 2h). Taken together, our findings suggest that the neurons analyzed, a majority of which were classified as distance cells, were in fact, grid cells.

**Fig. 2.**
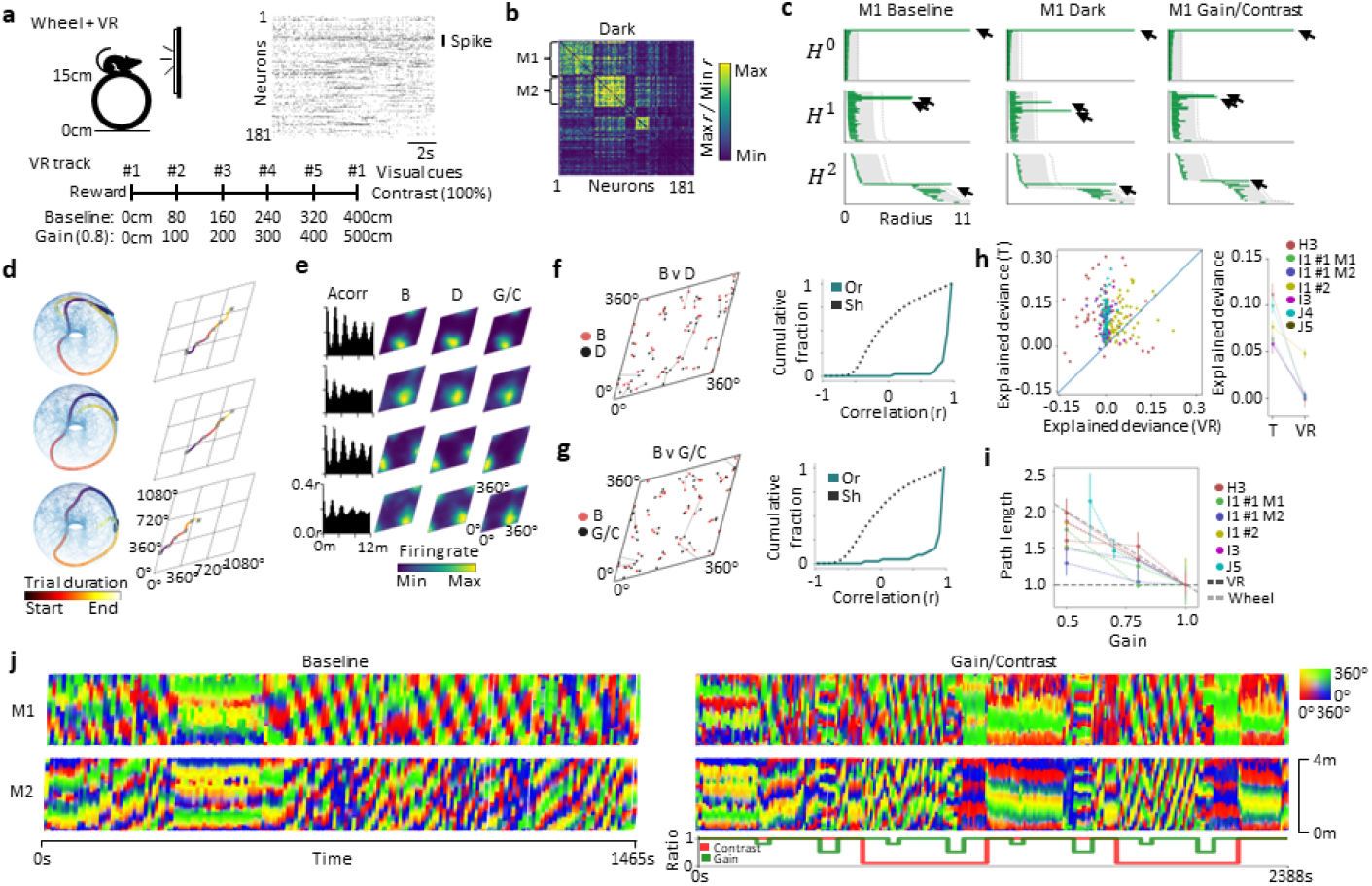
UMAPH detects tori in spiking activity of head-fixed mice during a VR task. **a**. Electrophysiological recordings were made while head-fixed mice engaged in a virtual reality task showing (for baseline trials) a continuously repeating linear VR track of 400 cm with five evenly spaced visual cues with a water reward at the end. During gain and contrast trials, the gain of the visual flow to the running wheel and the contrast of the visual cues were reduced. **b**. Cross-correlation matrix during dark session (monitors turned off and no rewards given) of mouse I1. Values indicated by color bar, minimum and maximum defined as 5th and 95th percentile. The two largest clusters, ‘M1’ and ‘M2’ (*n* = 44 and 41 neurons) are indicated. **c**. Barcodes (as in Fig. 1d, g) indicate a toroidal state space in every condition for M1. **d**. Example trajectories on the decoded neural torus for three different trials, embedded in 3-D (left) and linearized to 2-D (right). **e**. Virtual track autocorrelogram and toroidal firing rate maps (as in 1e and h) in baseline (B), dark (D) and gain/contrast (G/C) sessions for four example neurons of M1 (see Extended Data Fig. 8 – 11 for all neurons). **f, g**. Peak activity locations (left) for all neurons in M1 in B (red) vs. D and G/C sessions (black), and cumulative distribution of Pearson correlation between rate maps of M1 of corresponding conditions (blue) (as in Fig. 1i). **h**. Explained deviance scores (as in Fig. 1f) for T vs. VR positions, for GLM models fitted to spike train data during baseline trials. Right, mean explained deviance (*±*s.e.m.) per ensemble (*n* = 25 − 65). The toroidal description explained the data better than virtual track in each case (*P* ∈ [3.8 *·* 10^−9^, 7.5 *·* 10^−6^], *W* ∈ [307, 1747], Wilcoxon signed-rank test). **i**. Median and interquartile path length of the internal toroidal trajectory per trial relative to baseline trials of similar contrast (10 or 100%) for different gain values (*x* axis) (3 − 8 gain trials for each gain and contrast combination, and 44 − 178 baseline trials). Dashed lines indicate expected trial length if strictly aligned to distance on wheel (gray) or to VR track (black). **j**. Each VR-position (*y* axis) during B (left) and G/C (right) sessions for modules M1 (top) and M2 (bottom), color-coded by the instantaneous toroidal coordinates. Bottom right, time-dependent gain (green) and contrast (red) values, given as ratio of baseline.

We then clustered the remaining (88) MEC recording days and classified ensembles as grid cell modules if the cells’ toroidal tuning matched an idealized point source distribution on a hexagonal torus [52] (Extended Data Fig. 3b), as expected for grid cells [53] (Extended Data Fig. 3a)). We observed similar toroidal (grid cell) characteristics as for M1 and M2 in 6 more ensembles (Fig. 2h and Extended Data Fig. 2, 3b, c and 9 – 11).

### Alignments of the toroidal representation in VR are unstable but follow (gain/contrast) manipulations

Usually, decoding of the internal state space dynamics is linked through the tuning to an external, known covariate (e.g., spatial position, [22, 54]). By contrast, our method performs unsupervised inference of state space dynamics derived directly from the population activity, allowing unbiased comparison with external covariates. Coloring the 1-D VR trajectories with the 2-D toroidal coordinates, we observed a clear, time dependent relation between spatial and toroidal coordinates (Fig. 2j, Extended Data Fig. 4 and Video 3). At times, the movement on the torus was aligned with the spatial movement (visible in the figures as segments with a stable horizontal pattern), but would occasionally shift or drift, supporting findings by Low et al. [19]. For ensembles M1 and M2, this seemed to happen in coordination, in line with recent work [54]. However, gain and contrast manipulations clearly elicited a shift in the alignment between the torus and virtual space. When the contrast changed from high to low, an aligned representation quickly disappeared until the contrast was reset, upon which the mapping returned to the former representation. During strong gain manipulation (gain *<* 0.8 and contrast either 10 or 100%), the toroidal path length was longer than baseline sessions (*P <* 0.04, *Z >* 4 for six ensembles; Fig. 2i). This conjunctive influence of self-motion and visual cues is in line with Campbell et al. (2021).

## Discussion

By combining UMAP and PH, we obtained a powerful and noise-robust method that revealed toroidal topology of grid cell population activity in mice without reference to an external covariate, both in calcium imaging and with electrophysiology. Remarkably, this 2-D structure was found during head-fixation and 1-D wheel running, where it had so far not been observed… We observed a clear relation of the internal dynamics to movement, suggestive of path integration. While the toroidal structure was preserved across sessions, the alignment to the VR-track was affected by gain and contrast manipulations, and underwent shifts also during baseline trials.

Our results provide a proof-of-principle, demonstrating how a topological perspective on population coding allows us to study neural computations that go beyond the dimensionality of the task. This contributes to debates about whether simple, artificial stimuli are sufficient [e.g., in visual neuroscience 13, 14, 55] Thus, these results have the potential to shift the way we think about neural population coding, unlocking exciting possibilities for future studies of various neural systems using ideas from topology [32, 56]. As we have seen, the internal representation of grid cells can look distorted when mapped onto space, making spatial responses difficult to study. We propose grid cells and other cell types (e.g., head direction cells, see Extended Data Fig. 6) are best understood through their activity state space and not their external modulation.

By using an unsupervised state space discovery approach, we can study and compare population dynamics, not easily available at the single-cell level. This will be needed to understand recordings where the relevant covariates are of lower dimension than the neural representation, as in this study. Moreover, our approach is also useful when covariates are not known, for instance, in *cognitive spaces* [56], or where we have reason to assume higher-dimensional features in neural data (e.g., three- or higher-dimensional tori expected for conjunctive grid cells and combinations of modules [57, 58], Fig. 5c).

We look forward to future studies that embrace the toroidal (rather than 2-D spatial) nature of grid cells, which can be studied with a vast array of experimental methods in head-fixed 1-D and virtual environments [59, 60]. These simple task settings come with a number of benefits such as highthroughput experiments with large or movement-sensitive equipment, detailed animal tracking and permit trial-based, stereotyped analyses [11, 14]. We believe these findings open the path to new insights beyond what has been expected from such artificial settings [3, 4, 14], also in other brain regions and cognitive tasks [61, 62].

## Acknowledgments

We thank Zong et al., Obenhaus et al., Campbell et al. and Peyrache et al. for sharing data publicly and Per Kristian Hove for setting up databases. We furthermore thank the Department of Mathematical Sciences (NTNU) and the computing resources provided by IDUN. This work was supported by a grant by the Research Council of Norway (iMOD, NFR grant #325114).

## Author contributions

E.H., D.A.K. and B.A.D. conceptualized and proposed analyses. E.H. developed and performed the analyses. E.H., D.A.K. and B.A.D. interpreted data and results. All authors wrote the paper. B.A.D supervised the project and obtained funding.

## Supplementary Information

Not available for this paper.

## Author Contact Information

Correspondence should be addressed to E.H. or B.A.D.

## Methods

### Preprocessing of entorhinal recordings

Calcium events found in the ‘NAT.mat’-files of mouse 97045 days 20210305, 20210307, 20210308, 20210317 (renamed #1-4) and mouse 97046 days 20210308 (#1) and 20210312 (#2) were accessed from [63]. Duplicate cells listed in variable ‘RepeatCell’ of the corresponding ‘NeuronInformation.mat’-file were removed. As the neurons were recorded in two separate planes with a temporal offset, the activity was interpolated at similar frames using the package *‘scipy*.*stats*.*interp1d’*. Activity values less than 10^−10^ were subsequently set to 0 and neurons with average activity above 10 were excluded in the analysis. This resulted in 1-hour recordings of *n* = 228, 350, 293 and 428 (#1-4) neurons for mouse 97045 and 50- and 70-min. recordings of *n* = 635 and 637 (#1 and #2) neurons for mouse 97046. The data were speed-filtered at minimum 5 cm *s*^−1^, so to remove moments of inactivity. Population vectors with no activity were excluded.

PCA whitening was next applied to the data, to denoise and standardize the *n*-dimensional data (*n* being the number of neurons), projecting the *z* - scored population vectors to its *d* = 6 first principal components and dividing by the square root of the eigenvalues (see [64] for a thorough discussion of PCA as a preprocessing tool).

Due to computational complexity of computing barcodes, the size of the point cloud was reduced. First, a *radial* downsampling scheme was performed (as in [34]). The point with maximum absolute summed value was chosen as initial landmark point, and the Euclidean distance to the rest of the point cloud computed. Points closer than *E* = 0.5 were discarded and the remaining closest point to all sampled landmark(s) (defined as the maximum distance to all landmarks) was then picked. This process was iterated until exhaustion, vastly reducing the size of the point cloud (leaving approx. 30 − 50% of the points). While this method preserves the spread of the data, it is prone to keeping outliers. Thus, a second, density-based downsampling method was used (setting *κ* = 1000, see ‘Fuzzy Downsampling’), keeping *m* = 1000 points. Finally, UMAPH was applied to the reduced point cloud, using cosine metric, *k* = 1000 and ℤ _47_-coefficients (47 is chosen to not likely divide the torsion subgroup [49]).

The calcium events of Mouse 88529 were found in the ‘filtered spikes’ database table derived from the MySQL-dump file ‘dump.sql’ (available in [65]). The following sessions were used (and renamed): ‘419c1c6b319d0ddf’ (#1 W), ‘5b92b96313c3fc19’ (#1 OF), ‘d5a06b6a7630bb11’ (#2 W), ‘7e888f1d8eaab46b’ (#2 OF), ‘26fd0fbe1e205255’ (#3 W), ‘1f20835f09e28706’ (#3 OF), ‘59825ec5641c94b4’ (‘#4 OF’), ‘c43d9bd004db772b’ (#4 Obj1),’9190f2fccd52497e’ (#4 Obj2). The ‘filtered cells’-table was used to get cell IDs with a signal-to-noise ratio above 3.5 (see Supplementary Information in [51]). Grid modules were extracted by using agglomerative clustering, with average linkage, based on the ‘Manhattan’ distance between spatial autocorrelograms (in each recording day) found in the ‘grid score’-table (signal type ‘spikes’ and parameter set ‘B’). The largest cluster found, when setting a distance threshold of 2000 *r*, were analyzed, giving *n* = 126, 160, 113 and 162 neurons (recording days 1 – 4) with sessions lasting 18-24 minutes. The events were temporally convolved with a Gaussian kernel of *σ* = 0.27 ms and square rooted, before speed-filtering the time frames using a threshold of 10 cm *s*^−1^. The same pipeline as described above was subsequently applied, with *E* = 0.7 in recording days 1 – 3 and *E* = 0.55 in #4.

Neuropixels recordings in the MEC were performed by Campbell et al. of 20 mice in a total of 90 sessions and retrieved from [66]. Each spike time was replaced by a delta function (valued 1 at the time of firing; 0 otherwise) and temporally convolved with a Gaussian kernel with *σ* = 60 ms, before summing over all spike times (for each neuron), giving a continuous firing rate function. Firing rates were sampled every 10 ms and square rooted, before speed-filtering at minimum 10 cm *s*^−1^. Neurons not determined as ‘good’ (based on *“contamination, signal to noise ratio and firing rate”*, see Method details in [22]) in one recording or neurons with mean firing rates below 0.05 Hz or above 10 Hz were excluded. The remaining neurons were then clustered (see ‘Unsupervised clustering’), giving 65 clusters and a similar pipeline as described above was used, with *d* = 7, *E* = 0.9, *m* = 800 and *κ* = 800 for all ensembles.

### UMAPH pipeline

Given a dataset *X* ⊆ *ℝ*^*N*^, UMAPH first computes geodesic distances for a uniform manifold approximation of the data from which topological information is then studied using persistent homology. This approach was first used to find topological structure in grid cell population activity [10] and has similarities with concurrent work in [67], computing persistent homology based on geodesic distances.

Let *X* be a dataset and *d* : *X × X* →*ℝ* a dissimilarity measure of the points inherited from the ambient space.

1. Choose a number *k* and define for each *x* ∈ *X* its neighborhood set *N*_*x*_ = {*x*_1_, *x*_2_, …, *x*_*k*_}, such that for any *y* ∈ *X \ N*_*x*_, *d*(*x, y*) ≥ *d*(*x, x*_*k*_) and ordered by *d*(*x, x*_1_) ≤ *d*(*x, x*_2_) ≤ … ≤ *d*(*x, x*_*k*_). The extendedpseudo-metric, *d*_*x*_, is defined as

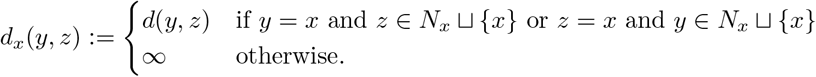

2. Next, for each *x*, define the (membership) function (or *fuzzy set*) *μ*_*x*_ : *X* → *I*, for *I* = [0, 1] by

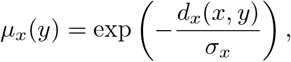

where *σ*_*x*_ is found by letting 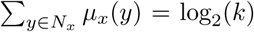. The value *μ*_*x*_(*y*) may be seen as the probability of the point *y* being a member of the neigbourhood of *x*.

3. Construct a global membership function *μ* : *X × X* → *I*:

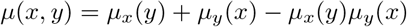

This operation is the *fuzzy* set union and defines the fuzzy topological representation of the data. This choice of union operation reflects the probabilistic sum, which describes the probability of the union of two independent events, and assigns probabilities to the edges of the neighbourhood graph of *X*. Other choices of fuzzy union operators are possible, and in [68] it is suggested that the fuzzy *intersection, μ*(*x, y*) = *μ*_*x*_(*y*)*μ*_*y*_(*x*), should be used for clustering. The standard fuzzy union is given by *μ*(*x, y*) = max(*μ*_*x*_(*y*), *μ*_*y*_(*x*)) [69] and is equivalent to taking the shortest path distance in merging the extended pseudometrics. This is used in [70] to describe the representation (or ‘UMAP complex’) as an iterated pushout of Vietoris-Rips systems.

4. Translate *μ* to a distance measure, *d*_*X*_ : *X × X* → *ℝ*,

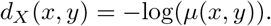

This converts the fuzzy topological representation back to an extendedpseudo-metric space representation of the dataset and *d*_*X*_ reflects the negative log likelihood of the simplices of the UMAP complex.

5. Finally, use *d*_*X*_ to construct a Rips filtration (see ‘Persistent Homology’) and take its homology. This may be seen as applying persistent homology to the filtration of the UMAP complex given by decreasing probability, giving a visualizable description of the fuzzy topological representation of the dataset *X* (i.e., the barcode).

*6. (Optional)* If the barcode indicates circular features in the data (captured by the one-dimensional bars), circular coordinates representing these may be computed through *co*homology. This requires defining the Rips complex for which the coordinates are computed, i.e., a choice of distance *τ* for which the circular feature exists, usually set to a value close to the end of the bar [71].

### Fuzzy Downsampling

To reduce computational complexity and remove outliers in the dataset, the point cloud was downsampled before applying persistent homology. Given a point cloud *X* and neighborhood sets *N*_*x*_, one for each *x* ∈ *X* (containing the *κ* closest neighbors), the global membership function *μ*(*x, y*) was computed as described in ‘UMAPH Pipeline’. Initializing *X*_0_ as an empty set, a subsample *X*_*N*_ ⊂ *X* was given recursively as follows.

For each iteration *n >* 0, define a function *F*_*n*_ as the summed membership strength for each point in the residual point cloud 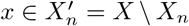, given by

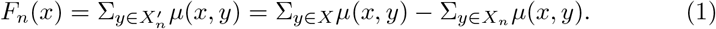

The (*n* + 1)-th point is then given by:

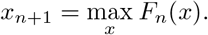

In other words, the method iteratively keeps the point with the highest probability of being in the neighborhoods of all other points.

Defining 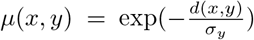, there is a close relation to mean-shift clustering [72] and the objective function used in the topological denoising technique introduced by Kloke and Carlsson [73]. The latter uses this function to translate a subsample of points to topologically relevant positions, describing it as a weighted difference of two Gaussian kernel density estimators, one for the dataset *X*, serving to push the subsample, *X*_*N*_, towards the densest regions of *X* and one for the subsample itself, repelling the points away from each other. Similarly, the fuzzy downsampling scheme picks points from the densest regions of *X*, but steers away from the regions already chosen.

### Persistent Homology

The shape of the neural data was characterized using persistent *co*homology. Persistent cohomology results in the same barcodes as persistent homology (which is described below), but cohomology was necessary for decoding [49]. The homology of a topological space, *T*, is a sequence of vector spaces *H*_*n*_(*T*), for all natural numbers *n* ∈ *ℕ* and the rank of *H*_*n*_(*T*) represents the number of *n*-dimensional holes [74]. A zero-dimensional hole describes a connected component, a 1-dimensional hole a circle, a 2-dimensional hole a void and so on for higher dimensions. The homology of a point cloud, *X*, only returns a count of the points. Thus, to elicit non-trivial homology reflecting the underlying space the dataset is sampled from, combinatorial spaces called *Rips complexes, T*_*τ*_ (*X*), are associated to the data. The Rips complexes depend on a scale *τ*, commonly describing a dissimilarity relation between the points in the point cloud. Varying *τ* gives rise to an ordered sequence of complexes known as the *Rips filtration*:

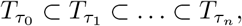

where *τ*_0_ *< τ*_1_ *<* … *< τ*_*n*_. Applying homology to the Rips filtration gives a sequence of vector spaces and maps induced by inclusion in each dimension, *n*:

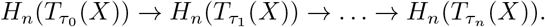

The totality of sequences is called persistent homology, *PH*_*_(*X*), and may be decomposed to a sum of *elementary persistence intervals, I*([*b*_*i*_, *d*_*i*_)):

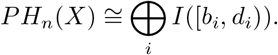

Here, *b*_*i*_ *< d*_*i*_ gives the scales for which an *n*-dimensional element hole in *PH*_*n*_(*X*) first appears and later disappears. Persistent homology may thus be represented as bars starting at *b*_*i*_ and ending at *d*_*i*_. The collection of such bars (across all dimensions) is called the *barcode*.

To obtain a shuffled distribution of the barcodes, the activity of each neuron was independently rolled a random amount of time. The same pipeline was then applied to the shuffled population activity to give a barcode, and the process was repeated 100 times with different seeds.

We used Ripser [75, 76] for all computations of persistent cohomology.

### Cohomological coordinatization

Circular coordinatization, as introduced by De Silva et al. [49], was performed to allow studying the internal dynamics of the population activity, This has previously been used to study head direction and grid cell activity [10, 34] and is motivated by a theoretical correspondence between 1-D cohomology and circle-valued maps of a topological space. By computing maps associated with the two longest-lived *H*^1^-bars in the barcode and the Rips complexes at *τ* = *b* + 0.99 *·* (*d* − *b*), where *b* and *d* correspond to the birth and death of the chosen bars, 2-D toroidal coordinates were computed for all vertices in the Rips complex. Note, for the head direction cell data only the single longest-lived bar was chosen.

The vertices correspond to the *m* points in the downsampled point cloud, thus, only *m* toroidal coordinates are obtained. To extrapolate these to the rest of the original point cloud or to a different session, the coordinates were first weighted by the values of the corresponding points, giving a distribution on the torus for each dimension. The toroidal coordinates were then computed, for each time step, by weighing the distribution by the corresponding value of each point in the full point cloud and finding the mass centers of the summed weighted distributions.

In visualizing the decoded toroidal positions as a function of VR-track (Fig. 2j and Extended Data Fig. 4), the coordinates were smoothed temporally with a Gaussian filter of *σ* = 0.5 s.

### Rate maps and autocorrelograms

Spatial positions in the open field were binned into a 30^2^ square grid, generating spatial rate maps. The mean neural activity in each bin was computed and spatially convolved with a Gaussian filter of width 2 bins (during which non-visited bins were assigned the mean value of the visited bins).

1D spatial autocorrelograms were computed for the linearized positions on the running wheel and the VR track, using 1 and 4 cm spatial bins, respectively. Autocorrelograms for each neuron were computed by finding the mean activity in each spatial bin and taking the dot product between this and a zero-padded copy of it, iteratively shifting the latter up to 300 bins.

Toroidal firing rate maps were calculated in the same way as the OF arena, first binning the toroidal surface into a square grid of 12° *×* 12° bins and computing the average activity in each position bin. To spatially smooth the toroidal rate map, the 60° angle of the toroidal axes was addressed by first shifting the bins horizontally by a length equal to half the bins’ vertical position. To address boundary conditions, nine copies of the shifted rate map were then tiled and spatially smoothed as the OF rate maps. The middle tile was then extracted and shifted back. For visualizations, the resulting rate maps were given 15° shear angles both horizontally and vertically.

### Unsupervised Clustering

The electrophysiological data were clustered into groups of neurons through the time-lag cross-correlation values between pairs of neurons. This was computed, as in [31], for the entire population recording:

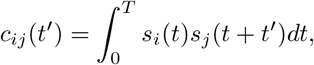

where *s*_*i*_(*t*) is the firing rate of neuron *i* at time *t*, converted from spike times as described in ‘Preprocessing of entorhinal recordings’, using a Gaussian kernel of *σ* = 0.3 s and sampled every 30 ms. *T* denotes the total duration of the recording. Setting *τ*_max_ = 0.9 s, the inverse, normalized cross-correlation was then given as:

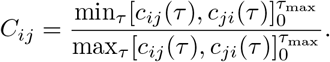

To cluster the neurons, we performed agglomerative clustering with average linkage on the squared *C*_*ij*_’s, averaged over all recordings of the same pair of neurons, using 0.6 *r* as distance threshold. Ensembles containing fewer than 20 neurons were disregarded, having too few neurons to confidently interpret the toroidal structure (see Extended Data Fig. 4e in [10]).

As described in the text, this was first performed for mouse I1, giving two ensembles, ‘M1’ and ‘M2’ (Fig. 6b). To see if these ensembles shared behavioral features, 1-D spatial autocorrelograms were computed. The mean autocorrelogram across the population showed repeating fields of activity along the track (Extended Data Fig. 3c). In both M1 and M2, the mean correlation between cells within the respective ensembles was higher than across the remaining MEC population in all three conditions (M1: 0.87 (B), 0.78 (D), 0.87 (G/C) v. 0.67, 0.58, 0.78; M2: 0.83, 0.82, 0.87 v. 0.71, 0.61, 0.77). Moreover, using the ‘distance cells’ classification in [22], 23 out of 44 (M1) and 27 out of 41 (M2) neurons were classified as ‘distance cells’.

### Hexagonal torus detection

To determine whether the decoded toroidal coordinates suggested a hexagonal torus in the VR-sessions, firing rates and rate maps were modeled for each neuron based on the analytical heat distribution on both a hexagonal and a square torus (Extended Data Fig. 3). The heat kernel on the hexagonal torus with point source at the origin is given as:

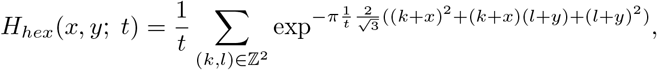

describing the temperature for (normalized) toroidal positions (*x, y*) ∈ [0, 1]^2^ after time *t*, while

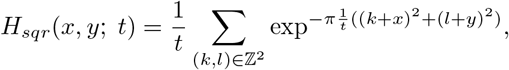

is the heat kernel on the square torus [52]. The fit of the original toroidal rate maps (generated by computing the mean activity in 12^2^ bins of the *m* toroidal coordinates found prior to extrapolation, see ‘Cohomological coordinatization’) for the VR data to a toroidal point source heat distribution was tested as follows. First, *m* toroidal coordinates were sampled with even spacing. The origin of the sampled torus was shifted to each cell’s peak activity on the torus (see ‘Comparison of toroidal tuning’), and using *t* = 0.1, *k, l* ∈ {− 1, 0, 1} in the above equations, allowed computing temperature estimates for each toroidal position. The heat distribution on the sampled torus was defined as the mean temperature in 12^2^ square bins. The linear correlation between the heat distribution and the firing rate maps asserted the hypothesized hexagonal toroidal tuning of each neuron. A hexagonal torus permits three periodic axes, and it was not known a priori pair of axes the decoded torus potentially described. However, this is reflected in the left vs right 45° angular shift of the distribution, so the reversed orientation was also tested by reversing one of the sampled coordinates. The maximum correlation between the modeled rate maps and the original one was used in the assessment and compared with a square torus rate map similarly modeled.

This analyses was first tested as a post hoc analysis of ensembles M1 and M2, showing that a majority of cells had rate maps more similar a distribution on the hexagonal torus than the square torus (M1: 38, 36, 36 out of 44 neurons in B, D, G/C and M2: 29, 35, 29 out of 41 neurons) and the median correlation was greater than by chance (M1 B, D, G/C: 0.81, 0.80, 0.75 *r* vs 0.08 – 0.16 *r* for 100 shuffles and M2: 0.71, 0.77, 0.68 *r* vs 0.02 – 0.11 *r*). The lowest median value for M1 and M2 was then used as a lower threshold to determine if the remaining ensembles had hexagonal toroidal structure, resulting in 16 such cases (6 mice and 8 ensembles, including M1 and M2, with *n* = 25, 37, 40, 41, 43, 44, 53 and 65 neurons for mouse and recording day: J4 0514, I5 0415, I1 0418, I1 0417 M2, J5 0506, I1 0417 M1, H3 0402, I3 0420. Recordings lasted between 9 and 159 minutes). Note, this heuristic eschews the computational problem of doing statistics on barcodes [77].

Replacing the sampled toroidal coordinates with the decoded toroidal positions in computing the heat kernel, *H*_*hex*_, gave time-varying heat models for the firing rate of each neuron. This allowed computing spatial autocorrelograms for idealized hexagonal toroidal tuning, visually matching the original firing rate counterpart (Extended Data Fig. 3c).

### Toroidal linearization and path length

The decoded coordinates were unwrapped onto a flat space to simplify the analysis of toroidal trajectories. This was done by iteratively assessing whether the next toroidal coordinate crossed either of the two circular origins (one for each dimension). All combinations (of 0 or 1 crossings per circle) were tested, and the next point was chosen to be the point closest to the previous one, as measured by the Euclidean metric.

To assess the influence of gain manipulation on the internal representation, the lengths of the linearized toroidal trajectories for each trial were estimated. First, the positions were fitted using linear regression for each axis and only trajectories with a fit of *r >* 0.5 in both axes were included. The length of each trial was then assessed as the Euclidean distance between the start and end point of the 2-D linear fit.

### Toroidal alignment

As the decoded origins and orientations of the toroidal descriptions are arbitrary, the decoded toroidal coordinates were first pairwise reoriented (without reference to space), in order to compare these across modules and sessions for the VR recordings. Moreover, it was necessary to account for the hexagonal torus allowing for three axes (each axis being a linear combination of the two other), with the decoded pair of axes oriented at either 60° or 120° relative to each other.

For each comparison, one set of toroidal coordinates was held fixed, with the goal of obtaining the same orientation and axes for a second set (i.e., session). First, in assessing the hexagonality of the decoded rate maps (see ‘Hexagonal Torus detection’), the angular orientation of the median rate map was determined for both sets of coordinates, limiting the possible axes-combinations (depending on the direction and if these were similar between the coordinate sets or not). To assess the remaining combinations, the coordinates of the fixed set were temporally smoothed with a 1-D Gaussian filter of width 200 ms and linearized. The time-varying directions of the toroidal trajectory was then computed as ‘arctan2’ of the derivatives of a cubic spline fitted to each unwrapped angle. The derivative of the sine and cosine of these directions were then compared with that of each combination of orientation and pair of axes for the second coordinate set. Note, in each assessment, the origins were aligned by subtracting the mean angular difference between the pair of coordinate sets. The combination that minimized the correlation between the two sets was recognized as the optimal alignment.

To align the toroidal coordinates of the ensembles with OF recordings, the combination of axes and orientation which looked most similar across different mice and sessions, when decoding the OF data and plotting the mean toroidal coordinate in 30^2^ square bins of the OF coordinates (Fig. 1a), was chosen.

### Generalized Linear Model

Bernoulli and Poisson generalized linear models were fitted to (binarized) calcium events and binned spike counts, respectively, using either the toroidal coordinates or the spatial positions as regressors, allowing a comparison of the explanatory power of the two descriptors. The setup was like Lederger et al. (2021) and Gardner et al. (2022), and included a smoothness prior to avoid overfitting.

Design matrices, *D*, were first constructed by binned the covariates and defining *D*_*i*_(*t*) = 1 if the covariate value at time *t* was contained in the *i* -th bin and *D*_*i*_(*t*) = 0 otherwise.

The probability of recording *n* spikes at time point *t* is given as:

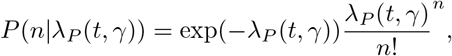

and the probability of recording a calcium event or not, *n* ∈ {0, 1 }, is given as:

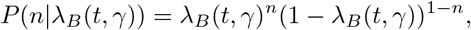

Here, *λ*_*P*_ (*t, γ*) = exp(∑_*i*_ *γ*_*i*_*D*_*i*_(*t*)) and *λ*_*B*_(*t, γ*) = exp(∑_*i*_ *γ*_*i*_*D*_*i*_(*t*))*/*(1 + exp (∑_*i*_ *γ*_*i*_*D*_*i*_(*t*))) is the expected activity at time *t* and the parameters *γ* were optimized by minimizing the cost function

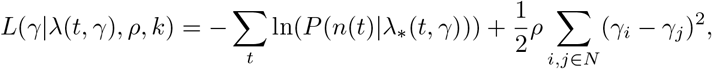

where *N* describes indices of neighboring bins. Note that the second term effectively minimizes large differences between neighboring parameters.

The minimization was run by first initializing each *γ* to zero and running two iterations of gradient descent on the above loss function before and after applying the ‘l-bfgs-b’-algorithm (given in the ‘scipy.optimize’-module with ‘gtol’=10^−5^).

The parameters *ρ* and number of bins were optimized for values 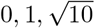 and 10 and 8^2^, 10^2^, 12^2^ and 15^2^, respectively for the OF arena of mouse 88592 day 2, and were found to be 1 and 10. These values were used for all recording days, and for mice 97045 and 97046. The 1-D VR-track positions were binned in 16 bins and *ρ* = 1 were used. To have the same number of bins for the 2-D toroidal positions, a 4 *×* 4 grid was used.

To avoid overfitting, the scheme was three-fold cross validated, i.e., the model was fitted to two-thirds of the data and its performance evaluated on the last third three times. The toroidal coordinates were decoded separately when used to fit each neuron’s firing rate, omitting the activity of the neuron studied in the extrapolation (see ‘Cohomological coordinatization’). Moreover, for the VR-sessions, baseline trials were decoded using the toroidal description found during dark session.

The deviance explained was computed, after similarly fitting a null model and a saturated model to the data, as:

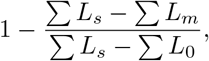

where *L*_*s*_, *L*_*m*_ and *L*_0_ are the log-likelihoods of the saturated, fitted and null model and the sums are over all time points in the given session.

### Comparison of toroidal tuning

Peak toroidal positions were visually compared across session to assess the conservation of toroidal tuning. The preferred locations were computed as the mass center of each neuron’s activity distribution on the torus (16^2^ bins), given by:

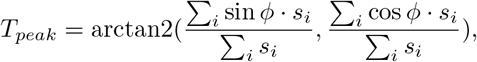

where *s*_*i*_ is the mean activity in the *i* -th toroidal bin *ϕ*_*i*_.

A second comparison was made by flattening the 2-D activity distributions (as above) and computing the Pearson correlation coefficient (using the ‘pearsonr’-function given in the ‘scipy.stats’-library) between two distributions from the same neuron in different conditions. To generate a random distribution, the cell indices in one session were randomly shuffled before computing correlation with the second session, i.e., computing the correlation of the rate maps of two different neurons. This process was repeated 1000 times.

### Continuous attractor network simulations

To study the toroidal structure of grid cell continuous attractor network (CAN) models, firing rate activity was simulated using three noiseless CAN models (Extended Data Fig. 3a).

First, a 56 *×* 44 grid cell network with purely inhibitory connectivity was simulated as proposed in [78]. Animal behavior was modeled by using the first 1000 s of the recorded trajectory of rat ‘R’ during OF foraging session day 1 found in [10]. The spatial positions were interpolated at 2 ms time steps and the speed *s*(*t*) and head direction *ϕ*(*t*) were computed as a function of the difference in position between each time step. The firing rate for time step *t*_*n*+1_ was given as:

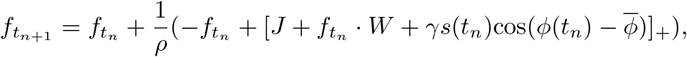

where […]_+_ is the Heaviside function, 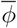 is the preferred head direction and parameters were given as: *J* = 1, *γ* = 0.15, *l* = 2, *R* = 20, *W*_0_ = − 0.01 and *ρ* = 10. The activity pattern was initialized to random and subsequently stabilized by running 2000 iterations of the above equation with no movement. The firing rate was set to 0 if below 0.0001 and resampled at 10 ms time steps.

Next, the twisted torus model by Guanella et al [53] was used to simulate a 20 *×* 20 grid cell network for a simulated random walk in a square box. The parameter definitions and code can be found in the open-source implementation by Santos Pata (https://github.com/DiogoSantosPata/gridcells). Here, we used 5000 time frames and ‘grid gain=0.06’.

Finally, an untwisted version of the previous model created square grid cell patterns for a 10 *×* 10 network. This was simulated using a Python translation of the Matlab implementation of Zilli [79]. 20 ms time steps were used, and a total duration of 295 s was simulated, following an OF trajectory recorded by Hafting et al. [1], provided in the same code repository.

### Head direction cell network

The UMAPH framework was applied to electrophysiological recordings from the mouse anterodorsal thalamic nucleus and subiculum of mouse 25 day 140130 (*n* = 25 neurons), mouse 17 day 130130 (*n* = 37 neurons) and mouse 28 day 140313 (*n* = 62 neurons) [80, 81]. The data contained 23to 192minute recordings during wakefulness, rapid-eye movement sleep (REM) and slow-wave sleep (SWS). Each mouse and brain state were processed and analyzed alike, similar to previously described in ‘Preprocessing of neural data’.

First, delta functions (one per spike) were convolved with a Gaussian kernel of *σ* = 250 ms for wake and REM sleep recordings and *σ* = 50 ms for SWS sleep. Population vectors were then sampled at 250 ms (wake and REM) and 2 s (SWS) intervals. Neurons were clustered as described in ‘Unsupervised clustering’ (based on the mean cross-correlation of the firing rates across brain states) and the largest cluster in each session was analyzed (*n* = 18, 25 and 41 neurons). In contrast to the entorhinal recordings, radial downsampling was not performed (i.e., *E* = 0), and parameters were set to *d* = 4, *m* = 500, *κ* = 300, resulting in barcodes indicating ring topology (Extended Data Fig. 6a). For visualization, the 6 first (whitened) principal components of the firing rates were projected to 3-D using UMAP (default parameters, Extended Data Fig. 6b).

Circular coordinatization of the longest-lived *H*^1^-bar was used to decode both awake and sleep data (Extended Data Fig. 6c). The decoded angles were reoriented by testing clockwise and anti-clockwise orientation and origin fixed by minimizing the mean difference to the recorded head direction angles during wake sessions. The angular tuning curves were computed using 60 bins and max-normalized.

### Three-torus

The applicability of the UMAPH framework in detecting higher-dimensional features was tested on a three-torus (*T* ^3^ = *S*^1^ *× S*^1^ *× S*^1^). First, 8^3^ = 512 points (i.e., three-dimensional angular values) were evenly sampled from a three-torus (Extended Data Fig. 5c) and added small, independent noise sampled from a normal distribution of width 0.1. The point cloud was embedded in ℝ ^6^ by taking the sine and cosine of each angle. Applying UMAPH with Euclidean metric and *κ* = 70 revealed the expected homology and the three circular features were decoded based on the three longest-lived *H*^1^-bars (see ‘Cohomological coordinatization’). To visually compare with UMAP, the 6-D embedding was projected to 3-D using UMAP (with 150 as number of neighbors).

### Data analysis and statistics

All data analyses were performed with custom-written scripts in Python 3.9. The following open-source Python packages were used: umap (version 0.5.2), ripser (0.6.1), numba (0.54.1), scipy (1.7.3), numpy (1.20.3), scikitlearn (0.24.2), matplotlib (3.4.2), h5py (3.6.0), gtda (0.5.1), cv2 (4.5.5), pandas (1.4.2) and datajoint (0.13.5).

The heaviest computational burdens were performed on resources provided by the NTNU IDUN/EPIC computing cluster [82] and that of the Department of Mathematical Sciences.

All statistical tests were one-sided.

## Code Availability

Code used in this article will be made available upon publication at GitHub.

## Data Availability

The data are publicly shared by the respective authors at: https://plus.figshare.com/articles/dataset/VR_Data_Neuropixel_supporting_Distance-tuned_neurons_drive_specialized_path_integration_calculations_in_medial_entorhinal_cortex_/15041316, https://archive.sigma2.no/pages/public/datasetDetail.jsf?id=10.11582/2022.00008, https://archive.sigma2.no/pages/public/datasetDetail.jsf?id=10.11582/2022.00005 and https://crcns.org/data-sets/thalamus/th-1.

**Video 1**. Comparison of open field foraging and toroidal population dynamics. Left, scatter plot of position in 2-D space of mouse 88592 day 2. Right, flattened torus, displaying toroidal population distribution at each time frame, given by first smoothing each neuron’s toroidal rate map (in 50^2^ bins) spatially with a Gaussian filter with *σ* = 3 bins. Next, every rate map was weighted by the firing rate of the corresponding neuron at each time point and the mean value for each bin across the entire population gave the time-dependent population distribution. Bright colors indicate high population activity.

**Video 2**. Comparison of wheel running and toroidal population dynamics for same neural ensemble as in Video 1. Left, wheel position (*y* axis) as a function of time (*x* axis). Right, as in Video 1 for internal toroidal representation decoded during wheel running.

**Video 3**. Comparison of VR running and toroidal population dynamics for gain sessions of mouse I3 recording day 0420. Left, wheel position as in Video 2 (top) and gain values (*y* axis) as a function of time (bottom). Right, inferred toroidal population dynamics, as in Video 1 and 2, found in the given recording.

---

**Extended Data Fig. 1.**
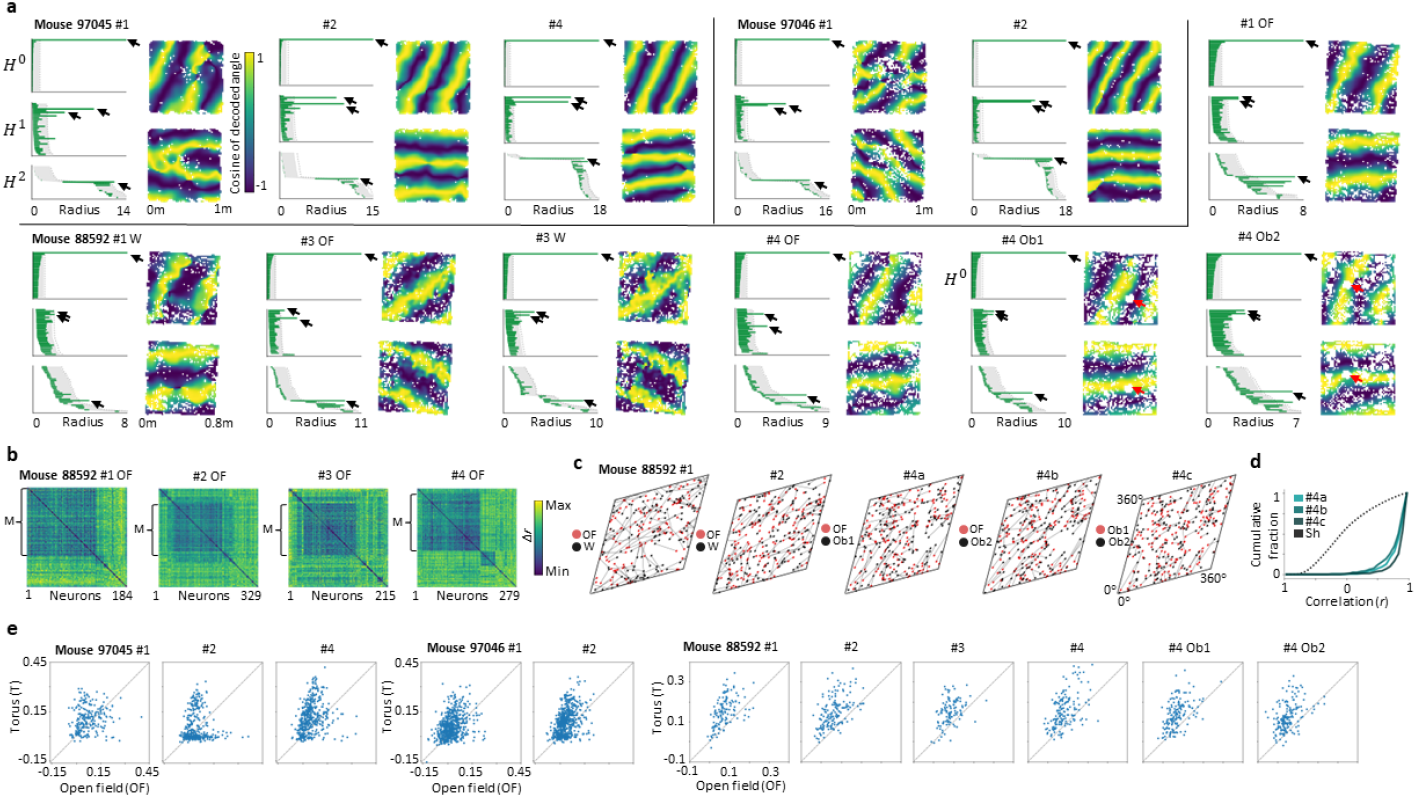
Barcodes, spatial mappings, distance matrices and toroidal tuning statistics for calcium recordings not shown in Fig. 1. **a**. Barcodes and mean toroidal coordinates as a function of OF locations (as in Fig. 1d) showing the results of UMAPH analysis for days 1 – 4 (*n* = 113 − 637 neurons). For wheel sessions, OF toroidal coordinates are found by using the longest-lived *H*^1^-bars to decode the activity during the OF session. The toroidal coordinates are reoriented to match across sessions. In two sessions (day 4 Obj1 and 2), an object was inserted into the environment (red arrows). **b**. Distance matrices of spatial autocorrelograms during OF foraging session for mouse 88592 across four different recording days. Values indicated by color bar. The clusters analyzed are marked as ‘M’ (*n* = 113 − 162 neurons). **c, d**. Distributions of receptive field centers and cumulative distributions of Pearson correlation of pairs of toroidal rate maps (as in Fig. 1i) for all neurons in each recording and comparison across environments. **e**. Comparison of explained deviance scores between OF locations and toroidal coordinates (as in Fig. 1f) for each neuron included in the UMAPH analyses in **a**.

**Extended Data Fig. 2.**
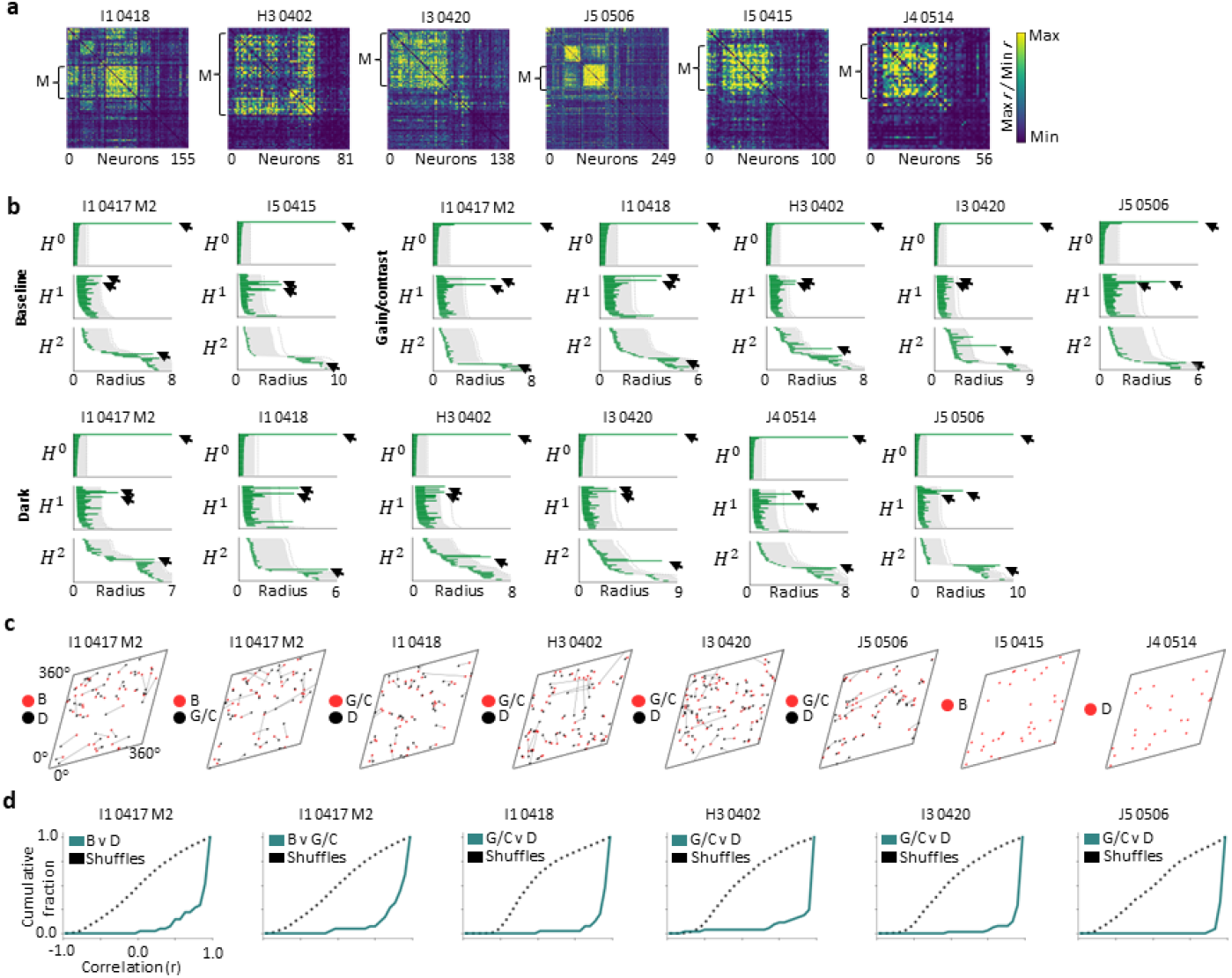
Correlation matrices, barcodes and toroidal tuning statistics for Neuropixels recordings classified as toroidal, not shown in Fig. 2. **a**. Correlation matrices (maximum divided by minimum) during dark session (as in Fig. 2b) sorted by cluster indices. Minimum and maximum range from 1.02 – 1.05 *r/r* and 1.59 – 2.86 *r/r* respectively. The clusters analyzed are marked as ‘M’ (*n* = 25 − 65 neurons). **b**. Barcode diagrams (as in Fig. 2c) for each session. **c, d**. Distributions of receptive field centers on inferred torus and cumulative distributions showing Pearson correlation of pairs of toroidal rate maps (as in Fig. 2f, g) for all neurons in each ensemble, compared across sessions.

**Extended Data Fig. 3.**
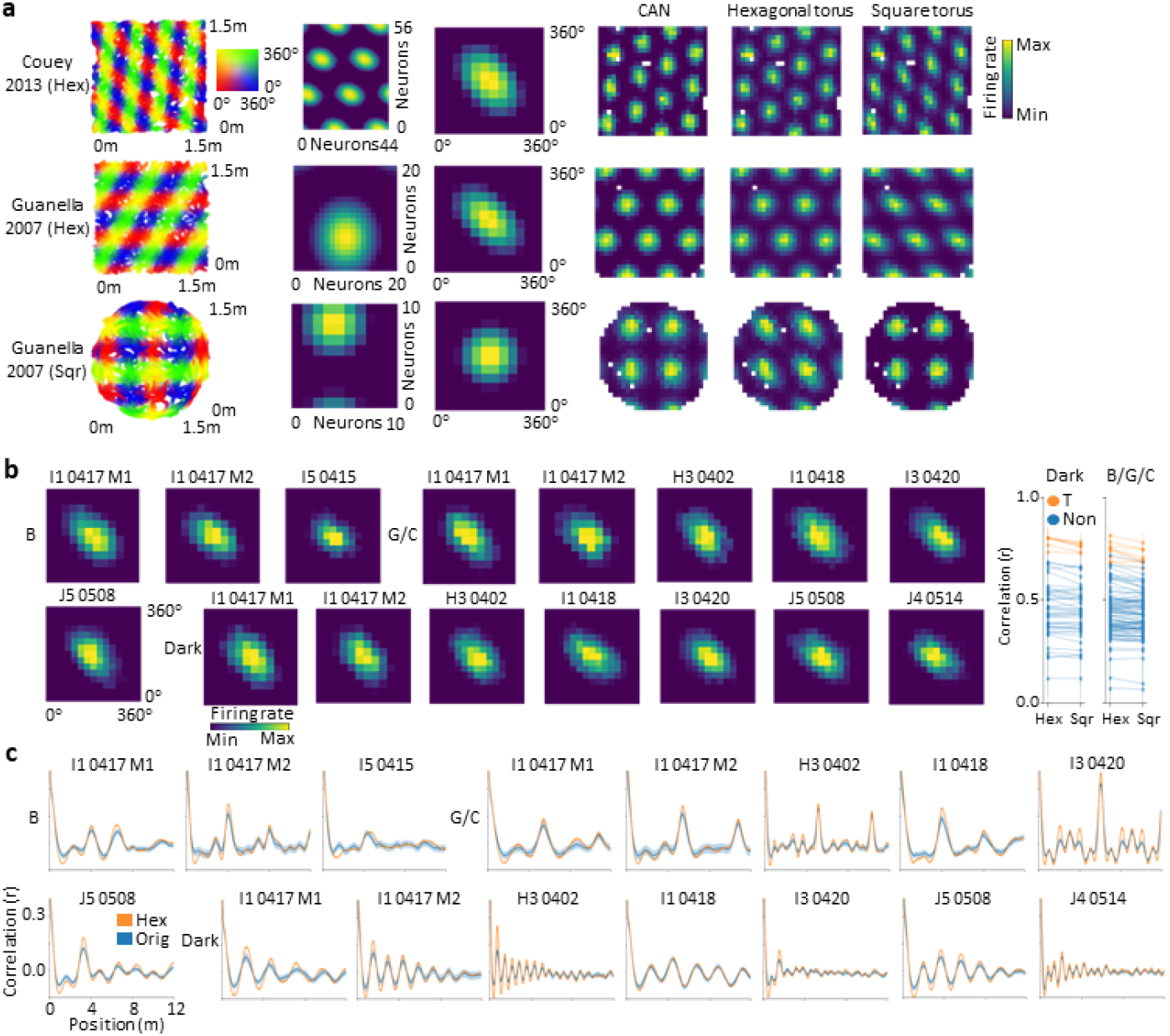
Decoding of toroidal coordinates in simulated grid cell and entorhinal ensemble activity allows comparing square and hexagonal toroidal structure. **a**. Circular coordinates from UMAPH analyses on data generated by three CAN models (row-wise, from top): an inhibitory, multi-bump network model (*n* = 2464 neurons), a twisted (*n* = 400) and an un-twisted (square, *n* = 100) torus network model. Each row displays (from left): a 2-D spatial trajectory colored according to toroidal position (2-D color map); firing rates of each neuron at a chosen time frame, ordered according to network connectivity (firing rate given by color bar); stacked, centered toroidal rate maps of all neurons in the network; spatial rate maps of activity from a CAN grid cell and fitted data using hexagonal and square torus point source models. **b**. Stacked, centered toroidal rate maps of all neurons in each ensemble (*n* = 25 − 65 neurons). Note angle of the firing fields. Right, median (and interquartile) correlation, per experimental condition, between each neuron’s rate map and the heat distribution on a hexagonal (Hex) and a square (Sqr) torus for all ensembles. Ensembles determined as toroidal are colored in orange. **c**. Mean spatial autocorrelograms (+- s.e.m) for each toroidal ensemble for recorded data (blue) and data generated by the hexagonal torus model (orange).

**Extended Data Fig. 4.**
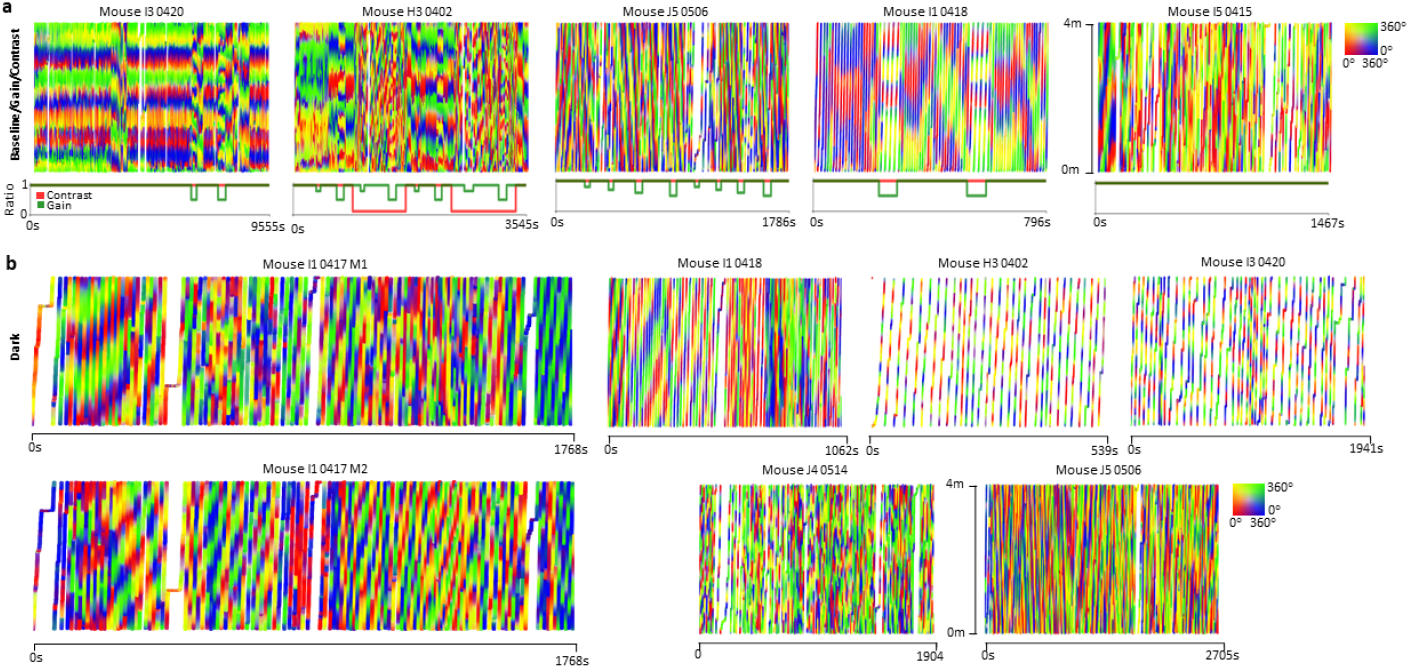
Internal toroidal dynamics as a function of the VR track for sessions not shown in Fig. 1j. **a, b**. Spatio-temporal positions colored by the 2-D toroidal positions across the entire session for baseline, gain, contrast and dark sessions. Gain and contrast manipulations are indicated in green and red line plots below relevant sessions. Note, in dark, the mice cannot see a progression in VR.

**Extended Data Fig. 5.**
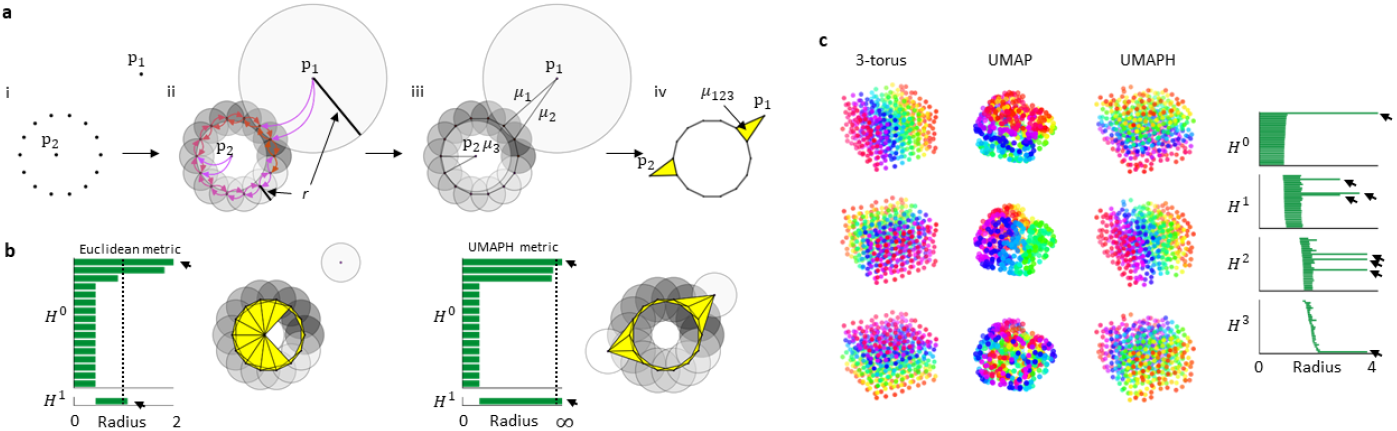
UMAPH visual description and example application for highdimensional manifold. **a**. The UMAP complex construction for a ‘noisy’ circular point cloud. i. Sample point cloud with two outliers, *p*_1_ and *p*_2_. ii. The geodesic distance of the underlying manifold is approximated by first computing local metric spaces (one for each point) through scaling the ambient space metric with respect to each point’s *κ* = 2 nearest neighbors. iii. The local metric spaces are merged to a global representation by translating each to a fuzzy set and then taking the fuzzy union of these. This creates a graph with weights *μ*_*i*_, describing edge (*i*) probabilities based on the scaled distance. iv. The UMAP complex is the weighted clique complex of this graph. I.e., the *n*-cliques form (*n* − 1)-simplices with weights equal to the minimum of that of its edges. Intuitively, *p*_1_ and *p*_2_ will lie similarly distanced from the circle. **b**. Left, applying PH to the point cloud in **a**, using the Euclidean metric, reveals a short-lived circle (*H*^1^-bar) and three long-lived clusters (*H*^0^-bars). Black arrows indicate the homology of a circle. The two additional clusters represent *p*_1_ and *p*_2_. The dashed line indicates the radius of the Rips complex shown to the right of the barcode. Right, the resulting barcode from applying PH to the UMAP complex (i.e., using −log *μ*_*i*_ as geodesic distances). The circle never closes, as seen in the corresponding Rips complex. **c**. Left, each column shows (from left): a noisy sample from a three-torus, *T* ^3^ = *S*^1^ *× S*^1^ *× S*^1^; 3-D UMAP projection of the 6-D embedding and 3-D embedding of the circular coordinates obtained from UMAPH. Each row is colored by the three sampled angles in the two first columns and by the decoded angles in the third. Right, barcode from applying UMAPH to the sampled 3-torus, with long bars implying the homology of a 3-torus (one *H*^0^-bar, three *H*^1^-bars, three *H*^2^-bars and one *H*^3^-bar, as indicated by arrows).

**Extended Data Fig. 6.**
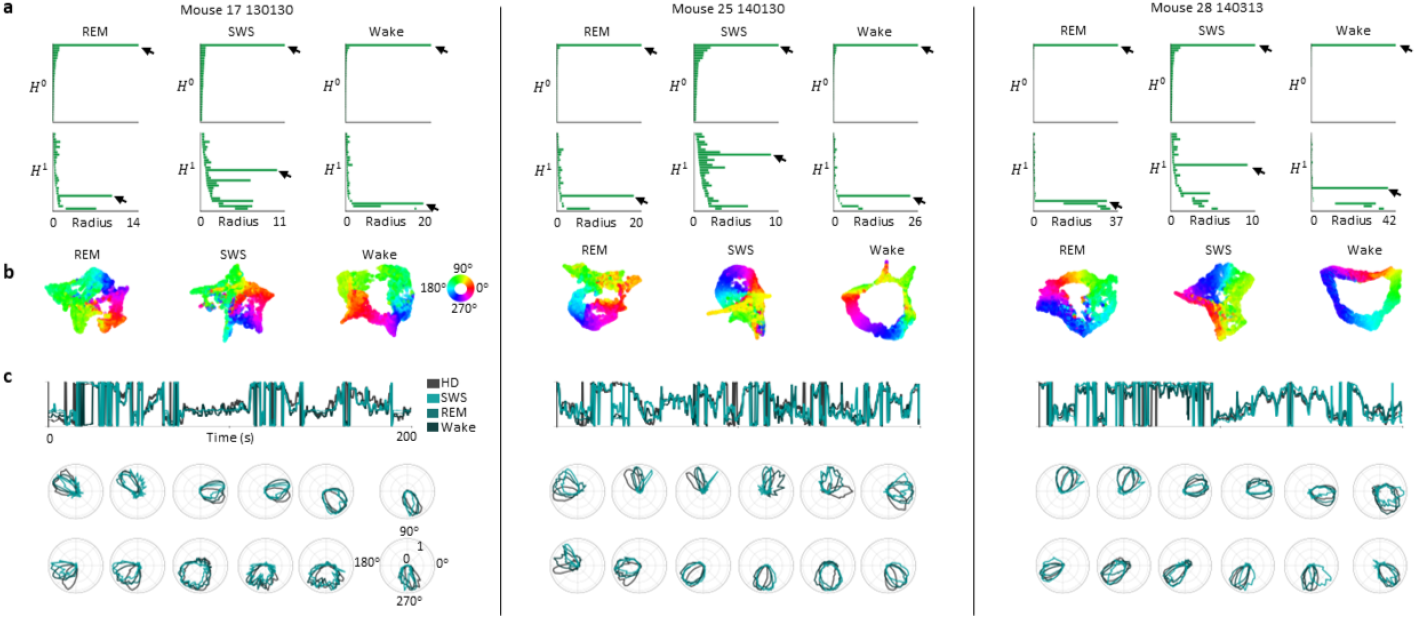
Ring topology of electrophysiological data in ADn and Subiculum representing internal head direction during OF foraging, REM sleep and SWS. **a**. Barcode diagrams from applying UMAPH in recordings of three mice (*n* = 18, 25 and 41 neurons). Arrows point to bars indicating ring topology. Each block (separated by black lines) contains the results from a single mouse. **b**. UMAP projection of firing rate activity colored by decoded circular coordinates (describing the circular features captured by the longest bar in the respective barcodes). **c**. Top, 200 s snippet of decoded circular coordinates and the recorded head direction. Note, the longest-lived circular feature found in each brain state was used to decode awake data. Bottom, single-cell tuning to the decoded coordinates and awake head direction recording for twelve example cells per animal (now, decoding each brain state separately).

**Extended Data Fig. 7.**
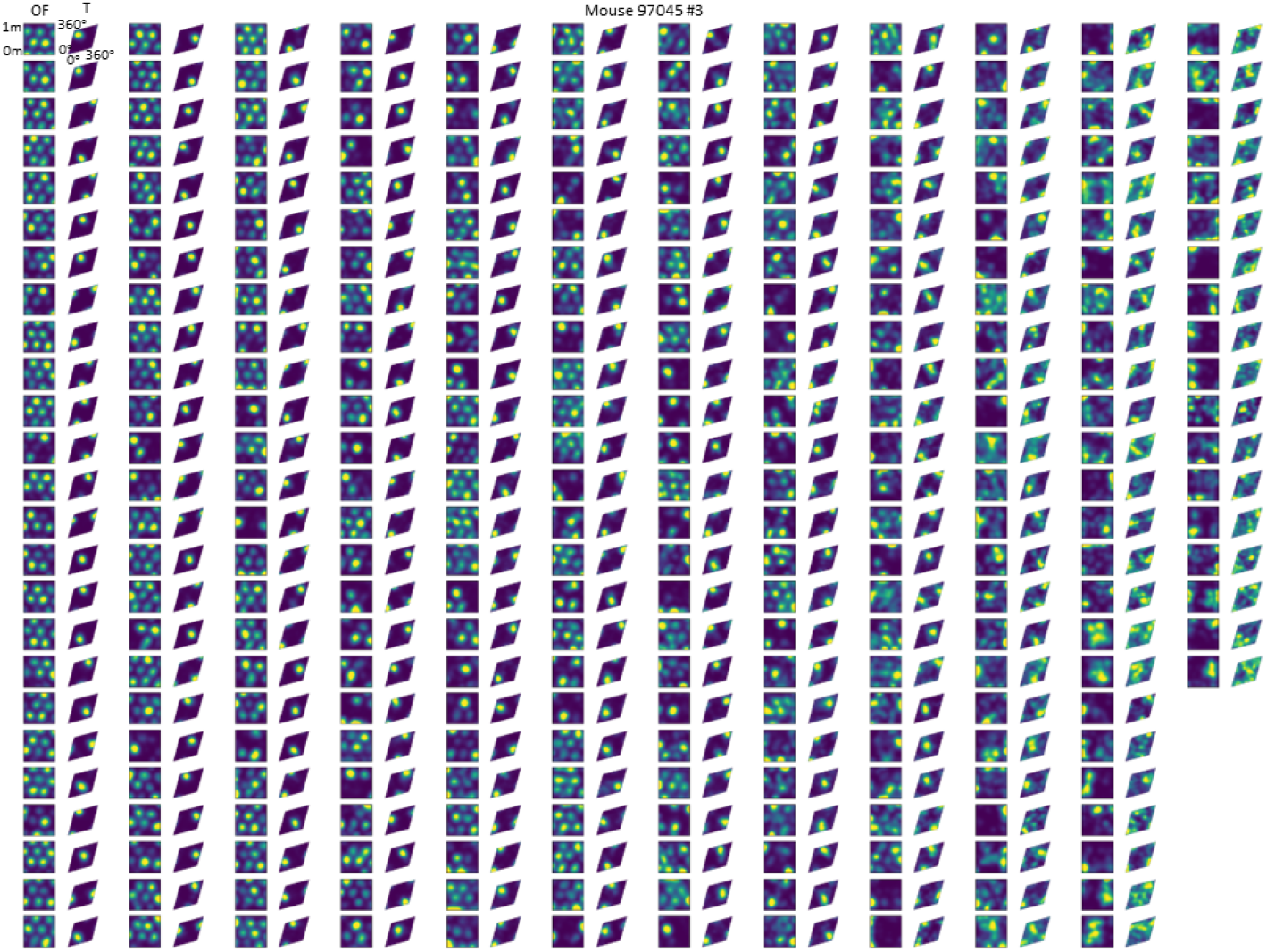
Tuning to coordinates in space and on the inferred torus for all neurons of mouse 97045 (as in Fig. 1e), recording day 2 (293 neurons) sorted top-bottom and left-right according to toroidal explained deviance (Fig. 1f).

**Extended Data Fig. 8.**
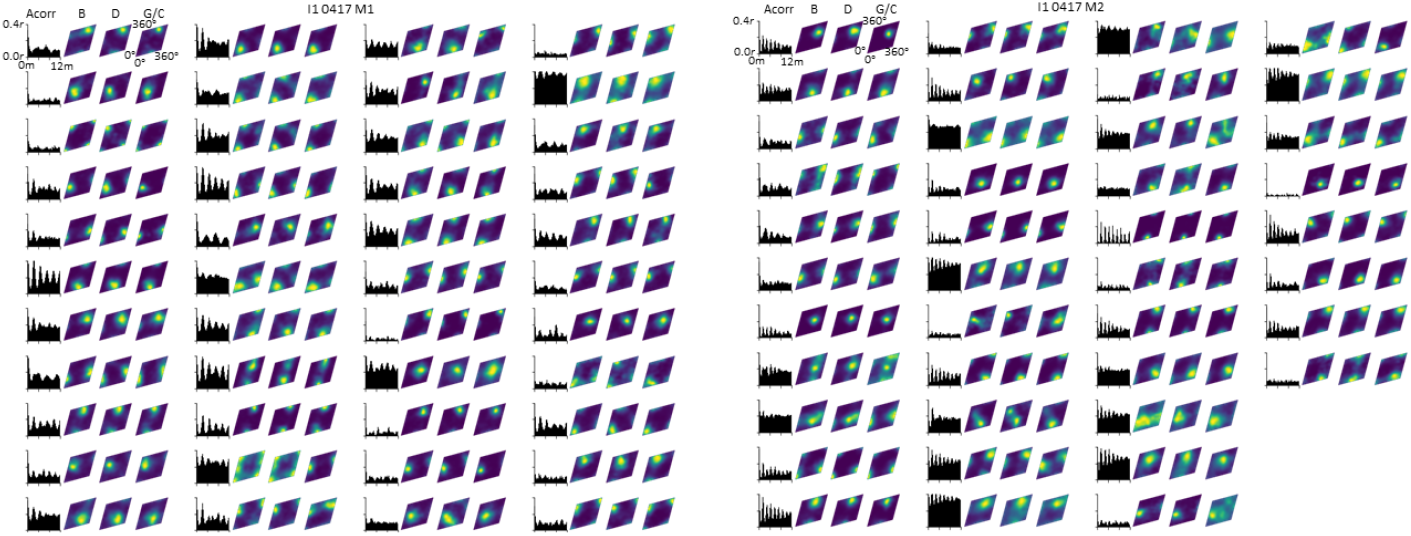
Tuning to coordinates in space and on the inferred torus (as in Fig. 2e) for all neurons of mouse and recording day (and module): I1 0417 (M1, 44 neurons) and (M2, 41 neurons). Each row of four shows 1-D VR-track autocorrelogram during dark session (left) and toroidal firing rate maps for each experimental condition (right).

**Extended Data Fig. 9.**
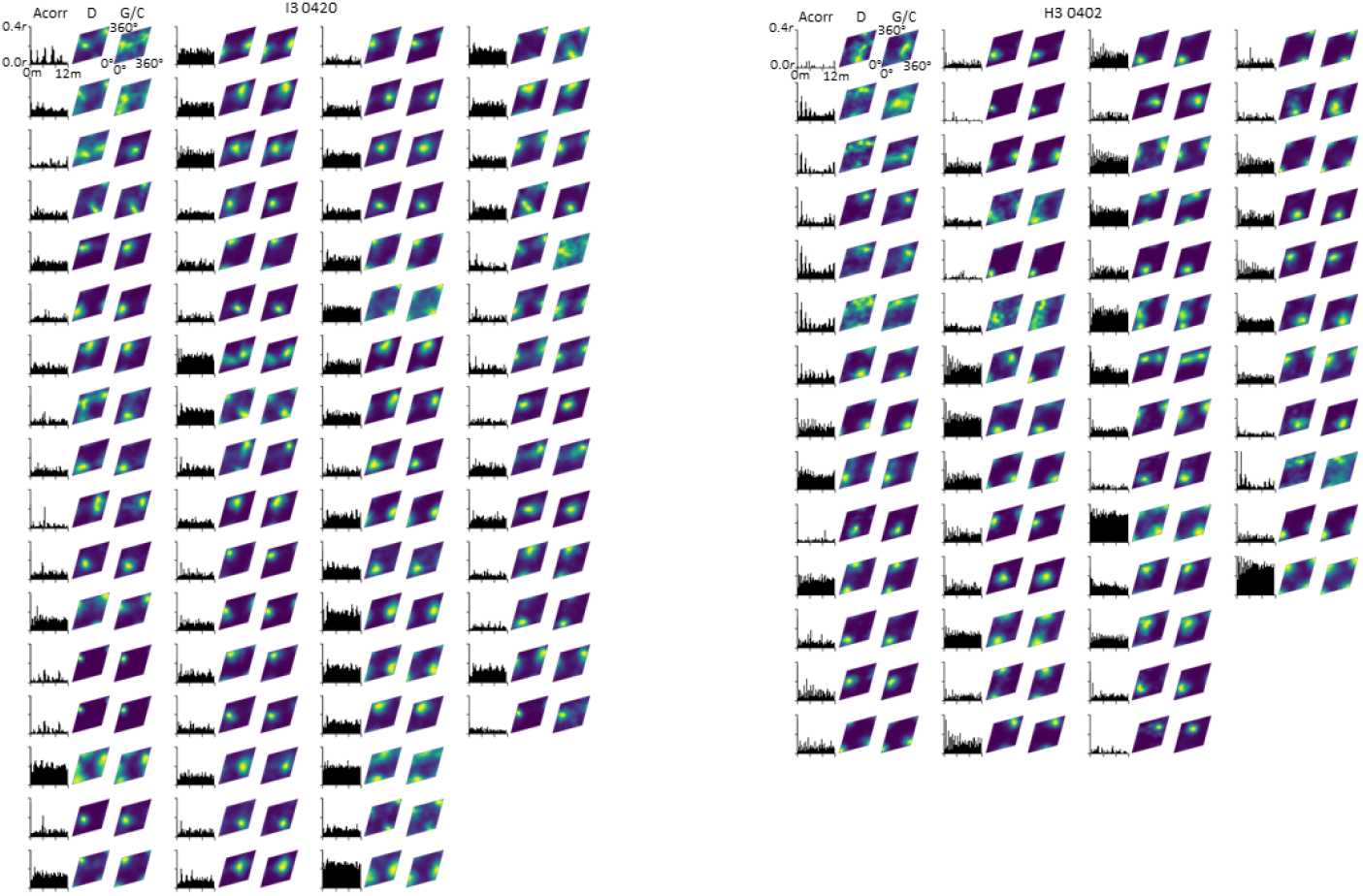
Tuning to coordinates in space and on the inferred torus for all neurons of mouse and experimental day: I3 0420 (65 neurons) and H3 0402 (53 neurons). Plots from left to right: 1-D VR-track autocorrelogram during dark session, toroidal firing rate map for dark (D) and gain/contrast (G/C) sessions.

**Extended Data Fig. 10.**
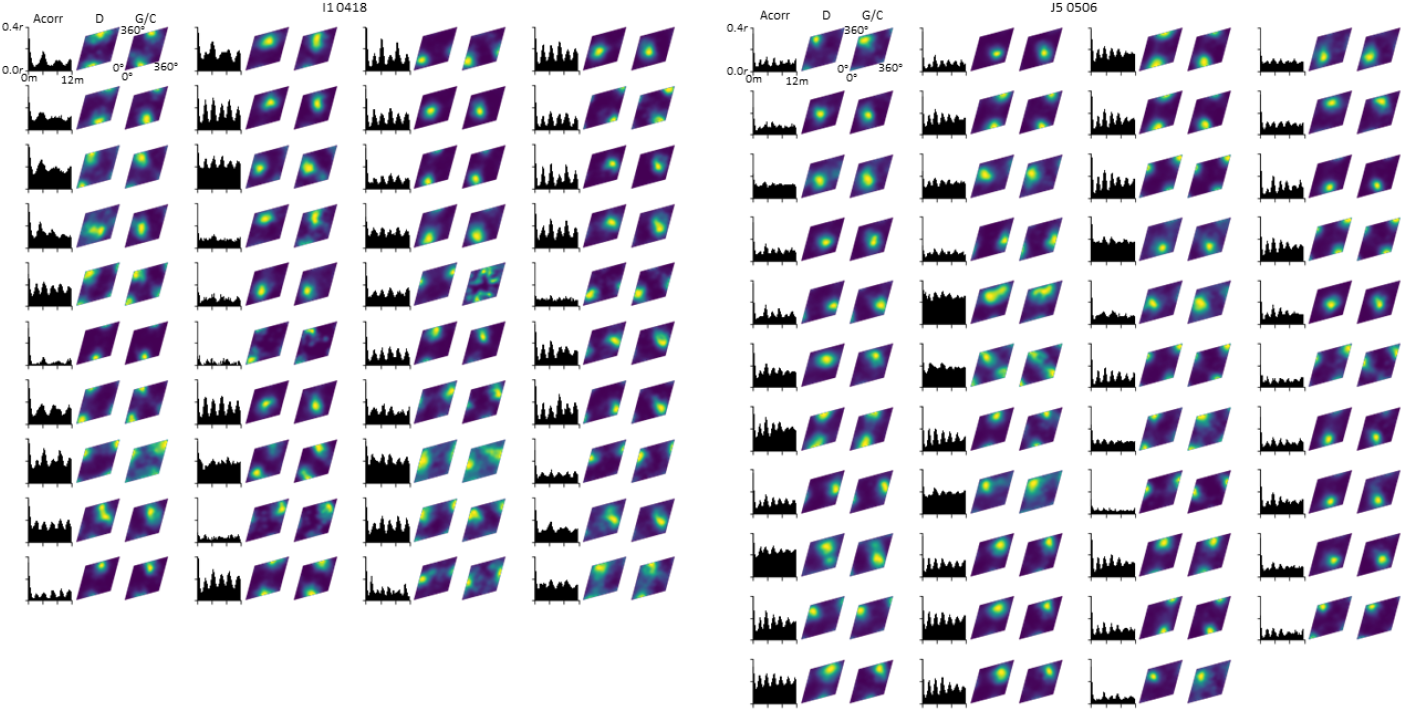
Tuning to coordinates in space and on the inferred torus for all neurons of mouse and experimental day: I1 0418 (40 neurons) and J5 0506 (47 neurons). Plots from left to right: 1-D VR-track autocorrelogram during dark session, toroidal firing rate map for dark (D) and gain/contrast (G/C) sessions.

**Extended Data Fig. 11.**
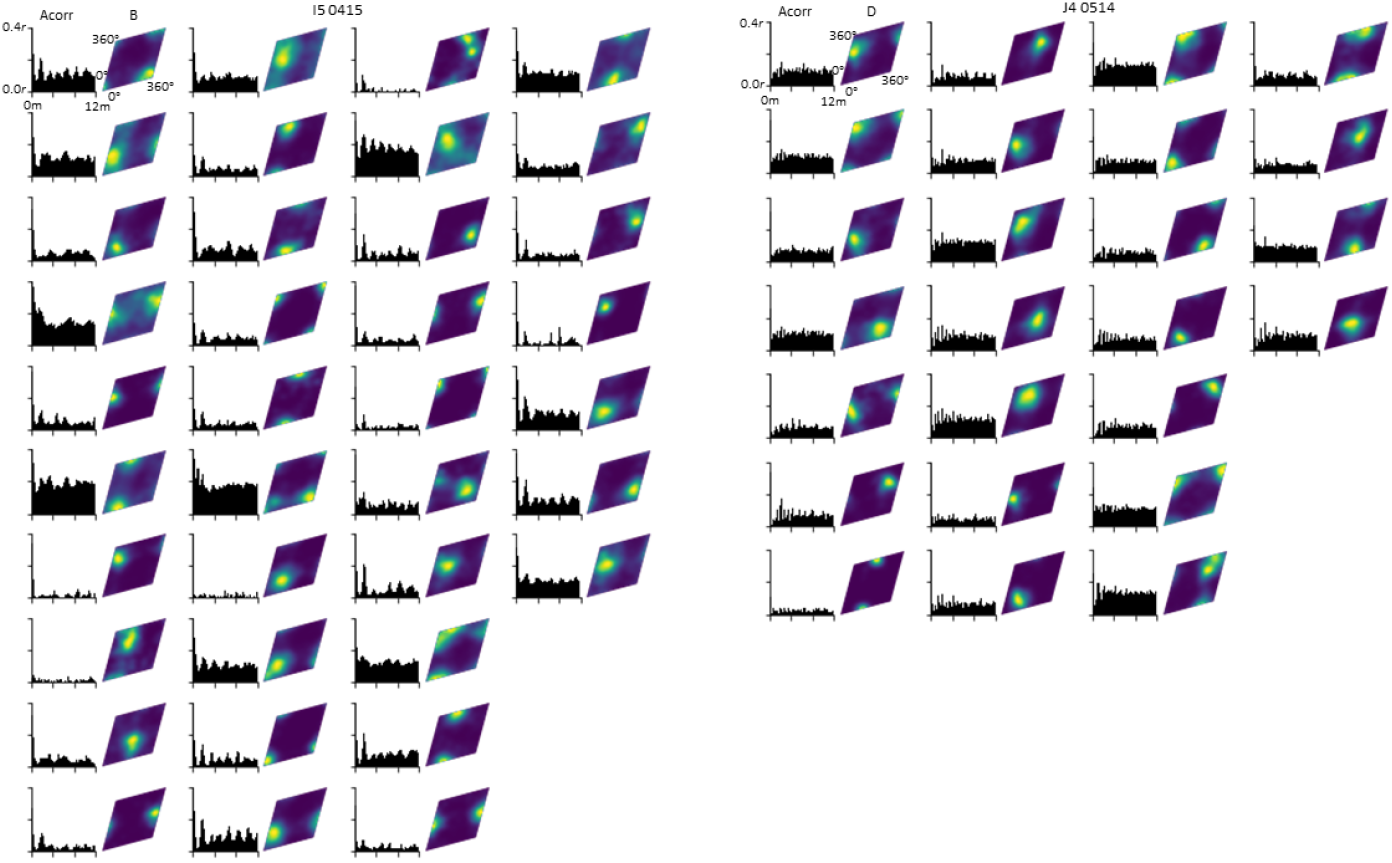
Tuning to coordinates in space and on the inferred torus for all neurons of mouse and experimental day: I5 0415 (37 neurons) and J4 0514 (25 neurons). Plots from left to right: 1-D VR-track autocorrelogram during dark session, toroidal firing rate map for baseline (B) or dark (D).

